# Glutamic acid promotes hair growth in mice

**DOI:** 10.1101/2020.09.27.315523

**Authors:** Carlos Poblete Jara, Beatriz de Andrade Berti, Natalia Ferreira Mendes, Daiane F. Engel, Ariane Maria Zanesco, Gabriela Freitas Souza, Lício Augusto Velloso, Eliana Pereira de Araujo

**Author notes:** Corresponding author: Eliana Pereira Araújo, Ph.D., Associate Professor, Faculty of Nursing, University of Campinas, UNICAMP, Tessalia Vieira de Camargo St, 126 - 13083-887 Campinas - SP - Brazil.

## Abstract

Glutamic Acid is the main excitatory neurotransmitter in neurons. Abnormal distributions of the glutamic acid receptors have been shown in hyper proliferative models such as psoriasis and skin regeneration. However, the biological function of glutamic acid in the skin remains unclear. Using *ex vivo, in vivo* and *in silico* approaches, we showed for the first time that exogenous glutamic acid promotes hair growth and keratinocyte proliferation. Topical application of glutamic acid decreased expression of genes related to apoptosis signaling in the skin. Also, we showed Glutamic acid increased viability and proliferation in cultured human keratinocyte. For the first time, we identified the excitotoxic GA concentration and we provided evidence for the existence of a novel skin signaling pathway mediated by a neurotransmitter controlling keratinocyte and hair follicle proliferation. In perspective, we anticipate our results could be the starting point to elucidate how exogenous glutamic acid from food intake or even endogenous GA from neuropsychiatric disorders modulate skin diseases.

## Introduction

Glutamic acid (GA) is the major excitatory neurotransmitter in the mammalian central nervous system and it is predominantly associated with excitatory synaptic neurotransmission (1). Previous reports have identified skin GA receptors and transporters across different species. Skin GA receptors subunits (Grin1, Grin2a, Gria2, and Grm1), and transporters (Slc1a1 and Slc1a2) were identified in rat skin (2), mouse skin (Grin1) and human keratinocyte (GRIN1) (3, 4)(5). Moreover, epidermis, hair follicles and sebaceous glands showed glutamate positive immunofluorescence (6).

Physiologically, the glutamatergic signaling through NMDARs in hair follicle cells was previously shown. Glutamic acid signaling is essential for innervation and differentiation of Grin1 positive Schwann cells during piloneural collar development in hair follicles (4). Specifically, NMDA receptors are highly expressed in type I and type II terminal Schwann cells. These cells are circumferential localized in the bulge border and covering a majority of outer root sheath keratinocytes in the isthmus (From Excitatory glutamate is essential for development and maintenance of the piloneural mechanoreceptor).

Previous reports have shown the effects of topical application of GA in wounded skins. *In vivo* studies showed that 1% L-Glutamic acid loaded hydrogels speeded up wound closure by increasing collagen synthesis and crosslinking in diabetic rats (7). Also, 1% L-Glutamic acid loaded hydrogels accelerated vascularization and macrophage recruitment into the diabetic wound (7). Moreover, topical D-Glutamic acid has also improved damaged skin by accelerating the barrier recovery (3), altogether suggesting a positive effect in skin repair.

We found five patent documentation using topical GA and derived molecules for hair growth stimulation, however, no scientific reports describing how GA could stimulate hair growth or epidermal cell proliferation. In this way, the biological effect of topical GA application in healthy skin is still unclear. Here, we evaluated the effect of topical GA in the back of healthy mice. We hypothesized whether GA could affect proliferation and viability in skin cells. In this manuscript, we have reported the preparation of L-glutamic acid formulations and its potential in hair stimulation.

## Result

### Glutamic acid increases human keratinocytes viability and proliferation

We hypothesized whether GA could stimulate proliferation and survival, keratinocytes will grow even in a confluence context (Fig. 1a). To do that, we removed any traces of FBS from the medium formulas due to the growth factors present in the bovine serum. By the end of 2 days of GA exposure, and even after 100% confluence, keratinocyte increased viability and proliferation. Indeed, we identified keratinocytes follow a Gaussian distribution pattern of viability after GA exposure (Fig. 1c). After 1 day of treatment, 100μM and 10mM GA concentrations increased human keratinocyte viability (Fig. 1b). These differences were higher after 2 days of GA exposure (Fig. 1c). The 100μM, 1mM, and 10mM GA concentrations increased keratinocyte viability within 2 days of treatment (Fig. 1c). As 10mM and 100mM GA concentrations showed opposite effects in the viability test (proliferative and excitotoxic respectively), we evaluated whether proliferation could be affected after 2 days of GA exposure (Fig. 1d). Consistently with viability results, BrdU positive keratinocytes were higher in the 10 mM group (Fig. 1g). On the other hand, we identified an excitotoxic concentration at 100mM GA. Keratinocytes treated with GA decreased cell viability after 1 day (Fig. 1b) and 4 days of treatment (Supplementary 1a).

**Figure 1.**
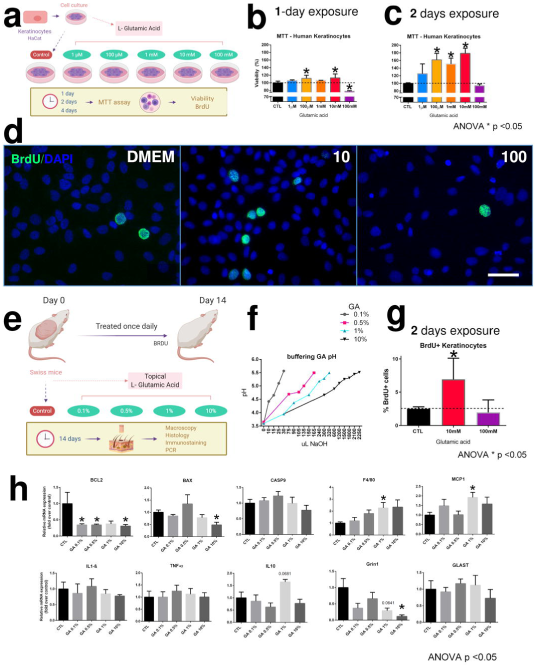
Effects of GA treatment in vitro and in vivo. Experimental design of cell culture experiment (a). HaCat cells viability results after 1 day (b) and 2 days (c) of exposure. Immunostaining of BrdU/Dapi positive cells (d). HaCat cells were exposed to DMEM (control group), GA 10 or 100 mM for 48 h and finally 3 h with BrdU, scale bar 50 μm. Proportion of BrdU-immunoreactive increased after exposure to GA 10 mM (g). Data is presented as mean ± SEM. N = 4 per group. p = 0.03 t-test Control vs Glutamic Acid 10mM; p = 0.03 One-way ANOVA. Experimental design of cell culture experiment (e). Different GA concentrations were equal to 5.5 pH (f). Bcl2, Bax, Casp9, F4/80, Mcp1, Il1β, Tnfα, Il10, Grin1, and Glast gene expression on skin samples after 14 days of GA treatment (h) * <p 0.05 ANOVA.

### Topical Glutamic acid decreases apoptotic related genes

To understand how GA promotes proliferation and improves viability, we evaluated apoptotic related gene expression. After 14 days of topical GA treatment, 0.1%, 0.5% and 10% decreased Blc2 gene expression (Fig 1h). Also, 10% GA treatment decreased BAX expression (Fig 1h). However, we found no differences in Casp9 expression (Fig. 1h). Next, we evaluate whether topical GA could stimulate expression of genes related to inflammatory response. We found no differences in Il1-β, Tnf-α, nor Il10. However, F4/80, a well-known marker of macrophage populations, and Monocyte Chemoattractant Protein-1 (Mcp1) gene expression were increased after 14 days of 1% GA (Fig. 1h). Additionally, topical GA 10% decreased the expression of Glutamate Ionotropic Receptor NMDA Type Subunit 1 (Grin1) with no differences in Glutamate Aspartate Transporter 1 (Glast) expression (Fig 1h).

### Topical Glutamic acid accelerates hair growth in healthy mice

To identify if our *in vitro* results could also be found in *in vivo* keratinocytes, we applied 4 different concentrations of GA on the back of Swiss mice (Fig. 1e, 2a). Surprisingly, 1% and 10% of GA speeded up hair growth after 14 days of topical treatment (Fig. 2a). Also, photomicrograph showed that GA increased external root sheath across all GA concentration (Fig. 2b, supplementary 1b) with no hyperkeratosis effect. Consistently, we also identified increased BrdU positive cells in the hair follicles and epidermal layer after 14-days GA topical treatment (Fig. 3a, d).

**Figure 2.**
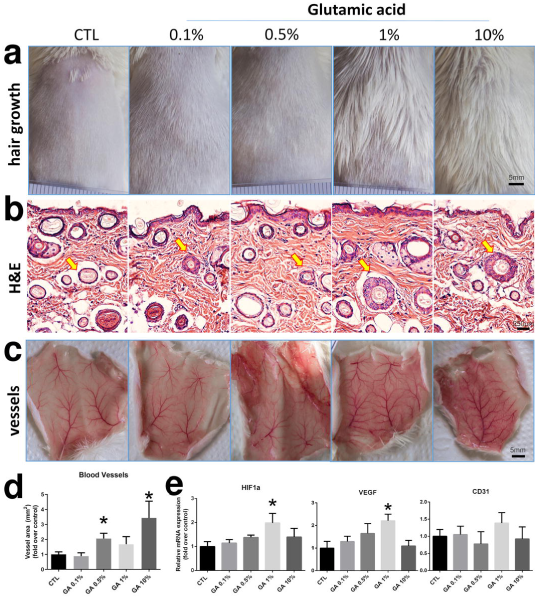
GA stimulates hair growth. Dose-response of topical GA application on the back of mice with Vaseline (CTL) and 0.1%, 0.5%, 1% and 10% GA for 14 days (a-c). Hair growth effect (a), H&E staining (outer root pointed with yellow arrows), and upside-down back skin samples showing vessel differences (c). Quantification of blood vessels area after 14 days of GA skin treatment (d). Gene expression of Hif1a, Vegf and Cd31 from back skin after 14 days of GA treated (e).

**Figure 3.**
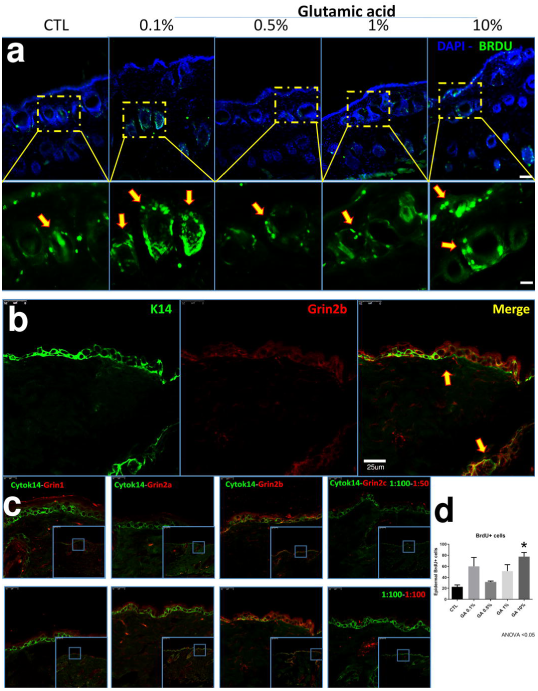
Topical GA increased BrdU+ cells. BrdU+ cells after 14 days of GA treatment (a). Yellow arrows indicate BrdU+ cells. Top scale bar 250μm and bottom 100μm (a). Characterization of different subunits Grin2b, Grin1, Grin2a, and Grin2c of GA receptor NMDA expressed in the skin of untreated mice (b-c). Yellow arrows indicate colocalization of K14+ Grin2b+ cells (b). Positive BrdU epidermal and hair follicle cells (d).

### Exogenous topical glutamic acid increased vascularization

We identified macroscopic differences in vascularization after 14 days treatment. The 0.5% and 10% GA topical application increased skin vascularization (Fig. 2c-d). Following these results, we evaluated whether GA could induce expression of vascular gene regulators. We found that 1% GA topical treatment increased the Hypoxia Inducible Factor 1 Subunit Alpha (Hif1a), a master regulator of vascularization (8, 9) (Fig. 2e). Also, 1% GA topical treatment increased the Vascular Endothelial Growth Factor A (Vegf), which induces proliferation and migration of vascular endothelial cells and it is essential for physiological angiogenesis (10-12)(Fig. 2e).

### Single cell RNA sequencing analysis showed differences in Glutamate receptor and transporter localization between mice and human

We evaluated glutamate receptor expression, using immunostaining, quantitative PCR, and single-cell RNA sequencing techniques. We identified that subunits Grin1, Grin2a, Grin2b, and Grin2c of NMDA receptor are expressed in the skin (Fig. 3b-c) and Grin2b is expressed specifically in keratin 14+ cells (Fig. 3b-c). To improve the accuracy, and due to the wide number of subunits (5 GA receptor families with 26 subunits) we used an RNA single cell approach (Fig. 4a). Using public transcriptome libraries of skin tissue, we analyzed ∼73,000 mice and human epidermal cells from back (mice), foreskin, trunk, and scalp (human). This cross-species analysis showed a similar percentage (5%) of “glutamatergic” epidermal subpopulation in the skin (Fig. 6b-c). In humans, we identified NMDA receptors as the highest expressed subunits in basal layer and hair follicular cell clusters, specifically GRIN2A subunit (Fig. 4b). Also, we identify that melanocytes expressed Glutamate Ionotropic Receptor Delta Type Subunit 1 (GRD1) (Fig. 4b). Additionally, we identified Excitatory Amino Acid Transporter 4 (SLC1A6) expression in granular cells, and Excitatory Amino Acid Transporter 1 (SLC1A3) expression in basal layer cells (Fig. 4c). In mice, we identified that Grin2d (in Sebaceous Gland) and Grik1 (in the hair follicle bulge) are the highest expressed subunits (Fig. 4d). Additionally, we identified the expression of Excitatory Amino Acid Transporter 3 (Slc1a3) (Fig. 4e) and Slc1a1 in 50% of all Sebaceous Gland cells (Fig. 4e)

**Figure 4.**
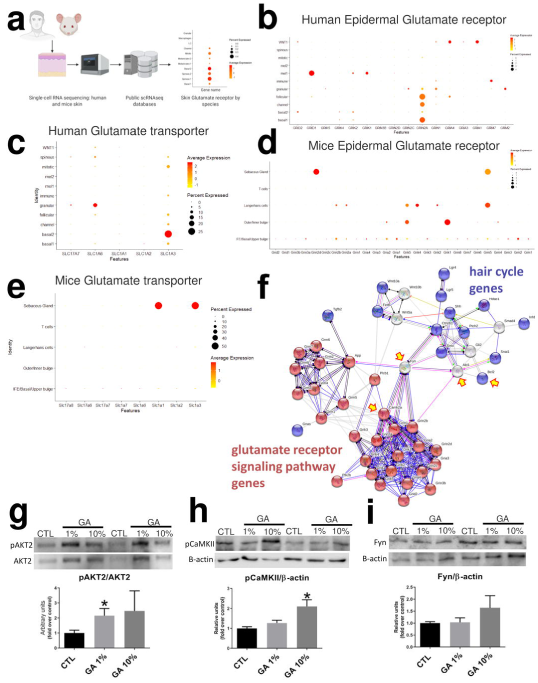
Cross-species skin GA receptor landscape using single-cell RNA sequencing. Generation Glutamic acid landscape using public data reveals GA distribution at single cell resolution in mice and humans. Schematic representation of our single-cell RNA sequencing analysis (a). Human Epidermal Glutamate receptors (b) and transporter expression (c). Mice Epidermal Glutamate receptors (d) and transporter expression (e). Glutamic acid and hair cycle Protein-Protein Interaction Network. Glutamic acid and hair cycle interactome were retrieved with the data-mining toolkit string. Interactors of Glutamic acid and hair cycle ontologies were set as colored nodes. Yellow arrows indicated shared shell interactors (f). Protein quantification to confirm Protein-Protein Interaction Network prediction (g-i). Phosphorylation of AKT (g), phospho-CaMKII (h) and Fyn quantification (i).

To predict a glutamic acid-mediated cell signaling pathway between GA signal and hair follicle related genes, we used computational interaction network analysis STRING. In this way, we used *glutamate receptor pathway genes* and *hair cycle genes* ontologies (Supplementary 2). We found that GA receptors interact with Hair cycle genes through the Tyrosine-protein kinase *Fyn*, Ca 2+ /calmodulin-dependent protein kinase II *(CaMKII)*, and Protein kinase B (*Akt)* (Fig. 4f). Additionally, we found *Bcl2* as a common apoptotic regulator between both hair cycle and GA pathway (Fig. 4f). To confirm that, we evaluated the protein expression of Fyn, CaMKII and Akt in the 14-day topical GA treated mice (Fig. 4g-i). We found no differences in Fyn quantification (Fig. 4i). However, we confirmed that AKT2 phosphorylation increased after 14 days of topical 1% GA treatment (Fig. 4g). Also, pCaMKII increased after 14 days of topical 10% GA treatment (Fig. 4h).

## Discussion

Interestingly, we found no scientific reports describing GA treatment or even GA effect in hair growth or epidermal cell proliferation. However, we found five patent documentation using topical GA (or derived molecules) for hair growth. One of these patents described the use of GA as a hair conditioner (patent number CN106580722A, China) for “hair restoration” and alopecia prevention. Other patent documentation showed a Poly-Gamma-GA composition for preventing hair loss and promoting hair growth (KR20150110149A, Korea). Also, synthetic compounds of GA attached to minoxidil for keratinocyte growth and hair growth in humans (USOO58O1150A, USA). Furthermore, a possible 2 to 12% GA topical cream could be used for combating hair loss or alopecia in humans (FR2939038B1, France). Additionally, 42 different molecules derived from L-Glutamic Acid were described as hair growth promoters (PI9302024A, Brazil). However, these patent documentations do not show any cell signaling to explain mechanisms of hair growth stimulation after GA application. In this way, the clinical importance of skin glutamic acid signaling is highly desirable.

Here, we tried to understand more about GA functions in the skin. First, using an *in vitro* approach we challenged 100% confluence human keratinocytes (without FBS or growth factor supply) to continue growing. Our result showed that GA increases proliferation and viability in keratinocytes. In this way, GA could represent a remarkable cheaper and highly available alternative to supplement culture mediums. On the other hand, previous reports showed that MK-801, an antagonist of GA receptor (NMDA receptor), decreased proliferation in primary human keratinocytes (2). Moreover, MK-801 topical application prevented hyperplasia induced by acetone (3), suggesting an anti-proliferative effect.

Glutamic acid has potent neurotoxic effects. Elevated amounts of GA lead to neuronal death in a process described as *excitotoxicity (13-15)*. GA transporters are a potent GA uptake system, acting as a neuronal compensatory response for *excitotoxicity*. GA transporter showed to prevent disproportionate activation of GA receptors by constantly removing GA from the extracellular space *(16-18)*. For the first time, we identified the excitotoxic GA concentration. *In vivo*, 100mM GA decreased keratinocyte viability and topical GA decreased Bcl2 and Bax expression. Altogether, our results support the excitotoxicity effect of higher concentration of pH-neutralized GA in keratinocytes.

To understand the exogenous GA effect on the skin, we identified the Glutamic acid transporter landscape at single-cell resolution in human and mice skins (Fig. 6d-e). Additionally, we showed Slc1a3 expression using quantitative PCR, and similar Slc1a3 (Glast) expression after exogenous GA application (Fig. 2c). Future research could help to identify the role of glutamic acid induced excitotoxicity and GA uptake system in the skin.

In terms of receptors, different subpopulation of glutamatergic cells has been extensively described (19-22). In the skin, previous reports identified the localization of the GA receptors and transporters in epidermis from rats and mice, as well as in human keratinocyte (2, 3, 5). These studies showed similar cross-species characteristics: a smaller subpopulation of cells expressing receptors and transporters (2). Consistently with our findings, here we showed a small subpopulation of epidermal cells expressing GA receptors along skin with varying of intensity (Fig. 5a-b). Our results suggest that these *glutamatergic keratinocytes* are responsive to exogenous GA stimulation.

Previous reports showed that vascularization increases during anagen of the hair cycle and decreases during catagen and telogen phases. This angiogenesis process was spatially correlated with upregulation of VEGF (23). Also, the hypoxia-inducible factor (HIF) has shown to coordinate up-regulation of multiple genes controlling neovascularization, as Vegf (24). Here, we showed that after 14 days of GA topical treatment on the back skin of mice, Hif1a and Vegf expression increased with remarkable change in angiogenesis as previously shown (7).

A recent study supports that glutamic acid-mediated signaling could be involved in hair growth (25). The authors showed that glutamine, a similar molecule to glutamic acid, controls stem cell fate in the hair follicle. The capacity of the outer root sheath cells to return to the stem cell state requires suppression of a metabolic switch from glutamine metabolism and it is regulated by the mTORC2-Akt signaling axis (25). Similarly, our result showed that glutamic acid increases AKT phosphorylation and outer root sheath cells. In this way, our results further suggest that GA activates the hair cycle by stimulating the stem cells to differentiate into outer root sheath.

Altogether, the clinical relevance of our study is to open a possible complementary signaling mechanism for hair growth disorders, or even for aesthetic hair stimulation using topical GA.

We encourage further research to uncover the relationship between skin disorders and Glutamic acid. Also, our results could be the starting point to elucidate how Glutamic Acid from food intake or even from neuropsychiatric disorders could be associated with skin diseases.

## Conclusion

Glutamic acid receptors and transporters are present in the cell of the skin. Human keratinocytes cultured with GA can increase viability and proliferation. Also, topical GA decreases apoptotic related genes, accelerates hair growth in healthy mice, and increases vascularization.

## Material & Method

### Experimental Animals

8-week-old male Swiss mice (n=6) were obtained from Breeding Animal Center of University of Campinas. Animals were maintained under pathogen-free conditions in individual cages on 12-12 hours dark-light cycle, at 21-23C°. Mice received chow and water *ad libitum*. Mice were anesthetized with intraperitoneal injections (according to the body weight), using ketamine hydrochloride 80 mg/kg and xylazine chlorhydrate 8 mg/kg. Under general anesthesia animal hair was removed using mechanical razor and depilatory cream (Veet) on the back of each animal (1cm x 2.5cm). The back of all animals was carefully cleaned to remove any trace of Veet cream. Animal experiments were approved by The Animal Ethical Committee at the University of Campinas, Brazil (certificate of approval no. 4930-1/2018).

### Topical Glutamic acid treatment

The back of the mice was treated once daily using different concentrations of Glutamic acid. To uniform 200 μL of treatment, we used different syringes preloaded with Vaseline (control), 0.1%, 0.5%, 1% or 10% glutamic acid (supplementary 1c). We spread the treatment manually using gloves which were changing between one group and another. To avoid removal of treatment, mice used Elizabethan collars (8 of 14 days of treatment).

### Topical glutamic acid composition

We made five different formulations from 0% to 10% of GA. Table 1 shows the different composition of each treatment. The pH of the formulations was adjusted using aqueous NaOH until the desired pH 5.5 was achieved. This pH value was chosen to resemble the skin surface pH (Fig. 2b)

### Cell culture MTT and BrdU

Human keratinocyte lineage (HaCat) passage 27-30 were culture in 37C°, 7% CO2 incubator with DMEM medium supplemented with 5% FBS to 100% confluence in 6-well plates. We replaced culture medium 2-3 times a week. 3-(4,5-dimethylthiazol-2-yl)-2,5-diphenyltetrazolium bromide (MTT) assay was used to analyze cell viability as previously described. MTT solution was prepared in Krebs-HEPES buffer (10 mM HEPES, 1.2 mM MgCl2, 144 mM NaCl, 11 mM glucose, 2 mM CaCl2, and 5.9 mM KCl). After 100% confluence (6-well plates), HaCat were incubated with the different concentration of Glutamic acid in DMEM without FBS for 1, 2 and 4 days. After treatment, the medium was removed and MTT solution (0.5 mg/mL) was added to each well and incubated at 37 °C for 3 hrs. Then the solution was removed and 300 μL of DMSO was added and incubated in dark at 60 rpm shaker. The absorbance product was measured at a wavelength of 540nm in a microplate reader (Globomax). All cell culture experiments were in quadruplicates. BrdU experiments were performed as previously described (26). Briefly, to assess the effect of GA on cell proliferation, HaCat human keratinocytes were maintained in Dulbecco’s modified Eagle’s medium (DMEM, Gibco) containing 4.5 g/L glucose, 4 mM L-glutamine, 100 units/mL of penicillin, 100 μg/mL of streptomycin, and 10% fetal bovine serum (FBS). Incubation conditions were 37 °C in 5% CO2/humidified air. HaCat cells were plated on coverslips in 24-well plates (1×10^5^cells/well) and exposed to GA for 48 h (10 and 100 mM) in DMEM without FBS. After treatments, cells were incubated with BrdU (10 µM, Sigma) for 3 h, then fixed with 4% PFA in 0.1 M PBS for 10 min at RT. For BrdU staining, cells were washed with PBS, and DNA was denatured with 1N HCl for 1 h at RT. Cells were blocked for 1 h in blocking solution containing 10% goat serum and 0.2% Triton X-100 in PBS, followed by an incubation with a primary (rat anti-BrdU; 1:200; Ab6326); and secondary goat anti-rat FITC (1:200; sc2011) antibodies prepared in 3% goat serum/ 0.2% Triton X-100 in PBS, and incubated overnight and 2 h, respectively. The nuclei were labeled with DAPI and coverslips were mounted onto glass slides for microscope imaging. Images were captured on fluorescence microscopy (Olympus BX41). The results of BrdU immunopositivity cells represent the average of 3 coverslips per experimental replicate, where 3 fields were imaged per coverslip and averaged. The number of immunopositive cells was quantified per image using the ImageJ software and are expressed as percentage relative to total DAPI nuclei.

### Animal photo Documentation

Hair growth processes were photo documented using a D610 Nikon digital camera (Nikon Systems, Inc., Tokyo, Japan). To secure a similar distance from camera to treated skin site, we used a stand and the same person to take photos.

### Histology

After 14 days of treatment, tissues were harvested and fixed by immersion in formaldehyde overnight. Any traces of formaldehyde were removed by 3 washes of PBS 1x. The tissue were processed in alcohol at different concentrations (70%, 80%, 95%, and 100%), xylol, and paraffin, before being fixed in paraffin blocks, and sectioned at 5.0 μm. 3 to 5 sessions were placed on microscope slides pretreated with poly-L-lysine. To evaluate cell and extracellular matrix morphology, the skin sections were stained with hematoxylin and eosin (H&E). The sections were incubated with hematoxylin for 30 s, rinsed in water, incubated for 30 s with eosin, rinsed again in water, and dehydrated. The slides were mounted in Entellan® and then analyzed; digital images were captured under bright-field microscopy.

### Protein-Protein Interaction Networks

Protein functional interaction networks were performed using STRING v11. The default functional interaction network was configured to *evidence* meaning of network edges, *experiments*, and *databases* in active interaction sources. *Mus musculus* organism was visualized by known molecular action. We analyzed two *Biological Process* using Gene Ontology Term from Mouse Genome Informatics database. A permalink webpage of *Glutamate receptor* (GO:0007215) and *hair follicle* (GO:0042633) Gene ontologies Interaction network is accessible through https://version-11-0.string-db.org/cgi/network.pl?taskId=lKUAbEZgGkVu for selected genes and https://version-11-0.string-db.org/cgi/network.pl?networkId=7i1fMIP01xqT for all genes.

### Single-cell RNA sequencing data acquisition, filtering, and processing

*In silico* analyses were performed using a HP ENVY 17 Leap Motion SE NB PC notebook with 16GB RAM and four-cores Intel i7 processor. Sample expression matrices (mice and humans) were downloaded from Gene Expression Omnibus and European Genome-phenome Archive: GSE67602 and EGAS00001002927. Cells were filtered by their total number of reads, by their number of detected genes and by their mitochondrial percentage. For mice we used nFeature_RNA > 10 and< 6000, nCount_RNA > 100 and < 50000, percent.mt < 9.5 settings. For humans we used nFeature_RNA > 100 and < 5000, nCount_RNA > 100 and < 25000 & percent.mt < 6 settings. Samples were processed in Seurat v3.1.5 using the default Seurat workflow. For clustering and visualization, we used the default Seurat pipeline gold standard and dot plot visualization. Cluster names were annotated to cell types accordingly original articles of Cheng et al. and Joost et al.

### Western Blotting

For the immunoblot experiments, the tissues were homogenized in RIPA lysis buffer (150 mM NaCl, 50 mM Tris, 5 mM EDTA, 1% Triton X-100, 0.5% sodium deoxycholate, 0,1% sodium dodecyl sulfate, and supplemented with protease inhibitors). Insoluble material was removed by centrifugation (11,000 rpm) for 40 min at 4 °C, and the supernatant was used for protein quantification by the biuret reagent protein assay. Laemmli buffer (0.5 M Tris, 30% glycerol, 10% SDS, 0.6 M DTT, 0.012 bromophenol blue) was added to the samples. One hundred micrograms of proteins were separated by SDS-PAGE and transferred to nitrocellulose membranes (Bio-Rad) using a Trans-Blot SD Semi-Dry Transfer Cell (Bio-Rad) for 1 hour at 17 V (constant) in buffer containing methanol and SDS. Blots were blocked in a 5% skimmed milk powder solution in TBST (1× TBS and 0.1% Tween 20) for 2 h at RT, washed with TBST, and incubated with the primary antibodies for 24 h at 4 °C. The primary antibodies used were anti-pCaMKII (Abcam, ab32678) and anti-Fyn3 (Santa Cruz, sc-16). HRP-coupled secondary antibodies (1:5000, Thermo Scientific) were used for detection of the conjugate by chemiluminescence and visualization by exposure to an Image Quant LAS4000 (GE Healthcare, Life Sciences). Anti-β-actin (Abcam, ab8227) was used as a loading control. The intensities of the bands were digitally determined by densitometry, using Image J software (National Institutes of Health).

### Immunohistochemistry

Skin expressions of Grin1, Grin2a, Grin2b, and Grin2c were identified by immunohistochemical staining. Immunohistochemistry was performed using the skin samples (n = 5). Tissue samples were immersed in 4% formaldehyde overnight. Tissue samples were washed three times with PBS 1x, cryopreserved in sucrose 20% for 3 days and 40% for 1 week. Samples were then embedded in OCT and sectioned using a cryostat (Leica CM1860). The sections (20 μm) were immunostained with the following primary antibodies: Grin1 (1:100, sc1467), Grin2a (1:100, sc1468), Grin2b (1:100, sc1469), Grin2c (1:100, sc9057), BrdU (1:200, ab6326) and keratin 14 (1:100, sc53253). VECTASHIELD with DAPI was used as a mounting medium for nuclear visualization. Images were obtained using a confocal microscope (Leica TCS SP5 II). For the in vivo BrdU experiment, we treated the mice intraperitoneally as previously described. Brightly, we applied one single injection of BrdU (150 mg/kg in buffer citrate) 3 h before skin harvest.

### Real-time Quantitative Polymerase Chain Reaction (RT-qPCR)

The total RNA content was extracted from the tissue using TRIzol reagent (Invitrogen). For each sample, two micrograms of RNA were reverse transcribed to cDNA, according to the manufacturer’s instructions (High-Capacity cDNA Reverse Transcription Kit, Life Technologies). Gene expression analysis via RT-qPCR was performed using TaqMan Universal PCR Master Mix (7500 detection system, Applied Biosystems). Primers used were: Bcl2: Mm00477631_m1; Bax: Mm00432051_m1; Casp9: Mm00516563_m1; F4/80: Mm00802529_m1; Mcp1: Mm00441242_m1; Il1b: Mm00434228_m1; TNFa: Mm00443258_m1; Il10: Mm01288386_m1; Grin1: Mm00433790_m1; Glast: Mm00600697_m1; Hif1a: Mm00468869_m1; Vegf: Mm00437306_m1; and Cd31: Mm01242576_m1 (Thermofisher). Analyses were run using 4 μL (10 ng/μL) cDNA, 0.625 μL primer/probe solution, 1.625 μL H2O, and 6.25 μL 2X TaqMan Universal MasterMix. GAPDH (Mm99999915_g1) was employed as a reference gene. Gene expression was obtained using the SDS System 7500 software (Applied Biosystems).

## Acknowledgements

The authors are grateful to Marcio Alves da Cruz, Joseane Morari, Vanessa Bobbo, Erika Anne Roman and Gerson Ferraz for technical assistance. This study was supported in part by the Coordenação de Aperfeiçoamento de Pessoal de Nível Superior – Brasil (CAPES) – Finance Code 88882.434714/2019–01.

